# Function within Disorder: Small heat shock proteins use different functional regions to chaperone tau aggregation

**DOI:** 10.64898/2026.01.28.702414

**Authors:** Mia Cervantes, Maria K. Janowska, Lisa M. Tuttle, Abhinav Nath, Rachel E. Klevit

## Abstract

In numerous neurodegenerative diseases known collectively as tauopathies, the microtubule-associated protein tau forms fibrillar aggregates that are hallmarks of disease pathology. Tauopathies represent a substantial fraction of diseases associated with protein misfolding. Cellular chaperones known as small heat shock proteins (sHSPs) play a critical role in maintaining protein homeostasis by delaying the onset of protein aggregation. Two sHSPs, HSPB1 (Hsp27) and HSPB5 (*α*B-crystallin), are constitutively expressed in brain and neurons. Here, we show that HSPB1 and HSPB5 delay tau aggregation *in vitro* through distinct mechanisms dictated by their disordered N-terminal regions (NTRs). HSPB1 inhibits tau aggregation under normal cellular conditions, whereas HSPB5 displays activity towards tau when activated by stress conditions such as pH acidosis. Using chimeric HSPB1/HSPB5 constructs in which small NTR subregions are swapped, we identify functional regions within the NTRs that modulate chaperone function for tau. The functional regions contain known sites of phosphorylation, suggesting that they are also control points that respond to cellular stress conditions. Our findings support an emerging model in which specific functional motifs within disordered regions of sHSPs govern activity and client engagement under normal and stress conditions.

**Broader Audience:** In many neurodegenerative diseases, the microtubule-associated protein tau forms fibrillar aggregates in the brain. Small heat shock proteins (sHSP) help prevent such aggregation, but their mechanisms of action remain enigmatic. We show HSPB1 and HSBP5, two sHSPs that are abundant and co-localize with tau, delay the onset of tau aggregation through distinct mechanisms. Each relies on specific small regions within their disordered N-terminal domains whose accessibility can be regulated by stress conditions and post-translational modifications.

## Introduction

Cells maintain proteostasis through coordinated systems that recognize, stabilize, and clear misfolded or aggregation-prone proteins. Failure in these systems contributes to age-related pathologies, most notably neurodegenerative diseases such as Alzheimer’s disease (AD). AD is the most prevalent of a class of neurodegenerative diseases known collectively as tauopathies in which toxic species of the microtubule-associated protein tau accumulate in brain and neurons (Creekmore 2024).

Tau is essential for neuronal microtubule stability and dynamics (Drubin 1986). In tauopathies including AD and fronto-temporal dementia, tau becomes hyperphosphorylated, dissociates from microtubules, and can assemble into amyloid fibrils that are hallmarks of disease pathology (Goedert 1989; Lee 1991; Jeganathan 2008; Lee 2011; Fitzpatrick 2017). Due to its intrinsic disorder, tau defies traditional structural biology tools and drug development pipelines, making progress towards therapeutics that productively engage tau challenging. As a result, the molecular mechanisms underlying tau toxicity, and the cellular defenses that counter it, remain incompletely understood.

Protein homeostasis is safeguarded by a network of molecular chaperones that suppress irreversible aggregation (Haslbeck 2005; Hartl 2011). Small heat shock proteins (sHSPs) are a first line of defense in maintaining a healthy proteome by stabilizing partly unfolded or misfolded proteins (“clients”) without ATP hydrolysis (Treweek 2015; Delbecq 2019). Clients stabilized by sHSPs may refold spontaneously or may be transferred to ATP-dependent chaperones like Hsp70 (Gonçalves 2021; Reinle 2022). sHSPs are constitutively expressed in many human tissues (Mymrikov 2017) and are sensitive to cellular stress signals, such as oxidation, post-translational modifications, and acidosis. These cues trigger structural rearrangements that enhance chaperone activity (Banzet 1998; Hayes 2009; Rajagopal 2015), positioning sHSPs to function under both normal and stress conditions.

sHSPs’ ability to delay protein aggregation without requiring ATP hydrolysis makes them attractive therapeutic targets for protein aggregation diseases. Indeed, efforts have been made to design small molecules or peptides that can bind to sHSPs to controllably activate their chaperone effect (Makley 2021). However, progress has been limited by an incomplete understanding of both the structure and mechanisms of how sHSPs interact with clients and of the activation mechanisms. This gap stems from properties of sHSPs that make them challenging to traditional biochemical/biophysical techniques: they are >50% disordered and form ensembles of large polydisperse oligomers ranging from dimers to ∼40-mers (Peters 2024).

All sHSPs share a tripartite architecture comprising a disordered N-terminal region (NTR), a folded α-crystallin domain (ACD), and a disordered C-terminal region (CTR). ACDs form dimers that serve as building blocks for higher order oligomers via inter-subunit interactions with NTRs and CTRs (Delbecq 2015; Clouser 2019). Each ACD dimer presents three conserved surface grooves: a “Central” groove at the interface between two ACDs and two “Edge” grooves at opposing ends of a dimer. These grooves accommodate potential binding partners, including but not limited to short stretches of the NTRs and

CTRs of other subunits within an oligomer (Rauch 2017; Baughman 2017; Freilich 2018; Clouser 2019). Because multiple binding sequences can bind to each groove, oligomers contain more binding partners than binding sites, giving rise to heterogeneity in sHSP oligomers. This “**quasi-ordered**” organization sequesters disordered regions within oligomers and creates structural and dynamic plasticity that is thought to govern sHSP function (Klevit 2020).

Of the ten human sHSPs, HSPB1 (Hsp27) and HSPB5 (αB-crystallin) are particularly abundant in brain and neurons (Mymrikov 2017, Bartelt-Kirbach 2017). Both are upregulated in AD, colocalize with tau fibrils in tauopathy brains, and exert protective roles against tau-mediated toxicity (Nemes 2004; Shimura 2004; Bjorkdahl 2008; Sahara 2007; Abisambra 2010; Hampton 2020). Despite these shared properties, mechanisms by which HSPB1 and HSPB5 engage tau, and whether they do so through shared or different strategies remain unresolved.

Here, we compare chaperone activities of HSPB1 and HSPB5 toward tau aggregation and define features that give rise to that activity. We aimed to disentangle two features of sHSP oligomers that together govern chaperone function: 1) the **local structural organization** that encodes for discrete chaperone-active regions and 2) the **oligomerization of sHSPs** that mediates the accessibility of specific regions. We find that HSPB1 suppresses tau aggregation under both normal and stress conditions and that HSPB5 requires stress-induced activation, for example, by pH to become effective. To identify the source of this difference, we focused on the NTRs, which are essential for tau chaperoning activity in HSPB1 (Baughman 2020). Using chimeric proteins, we show that scaffolding of intrinsically disordered regions within oligomers dictates chaperone potency. Finer-scaled chimeras in which short subregions of the NTRs were swapped between the two sHSPs identified specific regions that mediate activity. Integrating aggregation assays, NTR accessibility by Hydrogen-Deuterium Exchange (HDX-MS), and oligomeric size distributions by mass photometry (MP), we demonstrate that specific NTR subregions encode client-specific chaperone function. Together, the results provide a mechanistic framework for how sHSP oligomers tune their responses to aggregation-prone clients such as tau.

## Results

### HSPB1 and HSPB5 delay in vitro aggregation of tau using their disordered N-terminal domains

HSPB1 can inhibit tau aggregation in a range of *in vitro* contexts (Freilich 2018; Baughman 2020). HSPB5 is also constitutively present in neurons and most other cells (Janowska 2019), so we tested whether it can chaperone tau. To assess the chaperone activity of HSPB1 and HSPB5 towards tau aggregation, we performed in vitro assays in which fibrils are detected by Thioflavin T fluorescence. Spontaneous aggregation of tau is slow and requires an inducer to start aggregation. A sulfated polysaccharide, heparin, has traditionally been used to trigger aggregation of tau (King 2000; Barghorn 2004; Ramachandran 2011). However, while heparin is produced in mast cells and basophils, it is not in neurons (Lindahl 2013; Xu 2014). We used polyphosphate, a polyanionic chain of phosphate groups, to induce tau aggregation. Polyphosphate is present in neuronal cytoplasm and its levels increase with stress (Gray 2015; Lempart 2019), thus providing a more realistic context for our *in vitro* studies. Under our assay conditions, polyphosphate induces tau aggregation faster than previously optimized conditions using heparin as the inducer (SI Figure 1A). Both HSPB1 and HSPB5 delay tau aggregation, whether induced by heparin or polyphosphate. HSPB1 is more effective than HSPB5 under non-stress (“basal”) conditions (pH 7.5, 37C°; Figure 1 left).

**Figure 1.**
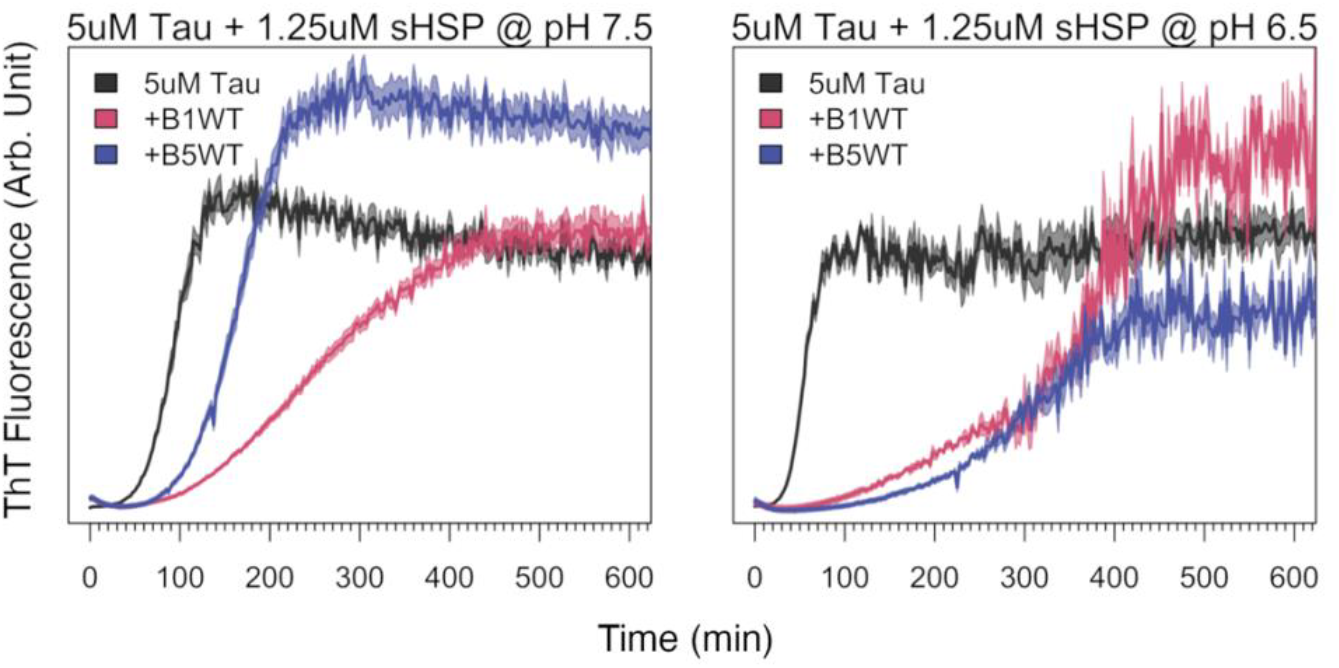
in vitro tau aggregation in the absence and presence of HSPB1 and HSPB5. Changes in Thioflavin T (ThT) dye fluorescence is monitored over the time-course of tau aggregation in presence and absence of sub-stoichiometric concentrations of HSPB1 or HSPB5. HSPB1 and HSPB5 chaperone polyphosphate-induced aggregation of tau at pH 7.5 (Left); HSPB5 becomes activated at pH 6.5 (Right). Both reactions are induced by 1 mg/mL polyphosphate using 5 μM Tau in presence and absence of 1.25 μM sHSP at 37 °C with agitation of the plate prior to each measurement. ThT fluorescence was recorded using excitation at 440 nm and 485 nm emission (arbitrary units). Solid lines represent the average ThT fluorescence, with lighter shaded regions indicating the SEM from 4-6 replicates per condition.

HSPB1 and HSPB5 are sensitive to environmental conditions. Among these, pH-dependent activation is the best characterized and is associated with various stress conditions including those in aging neurons (Hagihara 2024). As shown in Figure 1A (right panel), both HSPB1 and HSPB5 effectively delay tau aggregation at pH 6.5, despite more rapid aggregation of tau alone under these conditions. Thus, HSPB1 and HSPB5 have similar chaperoning capacity towards tau, but HSPB5 activity is lower under basal conditions.

The disordered NTR is required for HSPB1’s chaperone activity towards tau (Baughman 2020). When tau fibril formation was assessed in the presence of constructs lacking the NTR, “ACD-only” constructs of HSPB1 and HSPB5 both showed only modest chaperone activity at very high concentrations. At equivalent concentrations to full-length counterparts, the activities were negligible (SI Figure 1B-C). Thus, both HSPB1 and HSPB5 require their NTRs to effectively chaperone tau.

### Interplay of ordered ACD and disordered NTR modulates HSPB1 and HSPB5 activity

Interactions between ACDs and NTRs within sHSP oligomers are thought to play a role in their mechanism of action. Surface grooves on ACD dimers scaffold multiple discrete regions within NTRs (Clouser 2019) through inter- and intra-molecular interactions that sequester certain regions and affect activity. Certain HSPB5 NTR regions are more exposed at pH 6.5 than at pH 7.5 (Woods 2025). Based on the differential tau activity observed between HSPB1 and HSPB5 at pH 7.5, we hypothesized that HSPB5-ACD is more effective at sequestering its chaperone-active regions than HSPB1 under this condition, thereby limiting its chaperone activity.

#### ACD swapping affects chaperone activity

To test this hypothesis, we created two chimeric proteins in which the ACDs were swapped: B151 (HSPB1 containing HSPB5-ACD) and B515 (HSPB5 containing HSPB1-ACD). Strikingly, ACD swapping has opposite effects on tau chaperone activity. B151 is a better chaperone at pH 7.5 than WT-HSPB1 and B515 is a less effective chaperone than WT-HSPB5 (Figure 2A). Quantitatively, the time required to reach 50% of the ThT signal at saturation (T_50_) increased by three hours for B151 compared to HSPB1, while B515 reached T_50_ 45 minutes earlier than HSPB5 (SI Figure 1D). These results suggest that the HSPB1 NTR is less effectively sequestered by the HSPB5 ACD and the HSPB5 NTR is likely better sequestered by the HSPB1 ACD of HSPB1.

**Figure 2.**
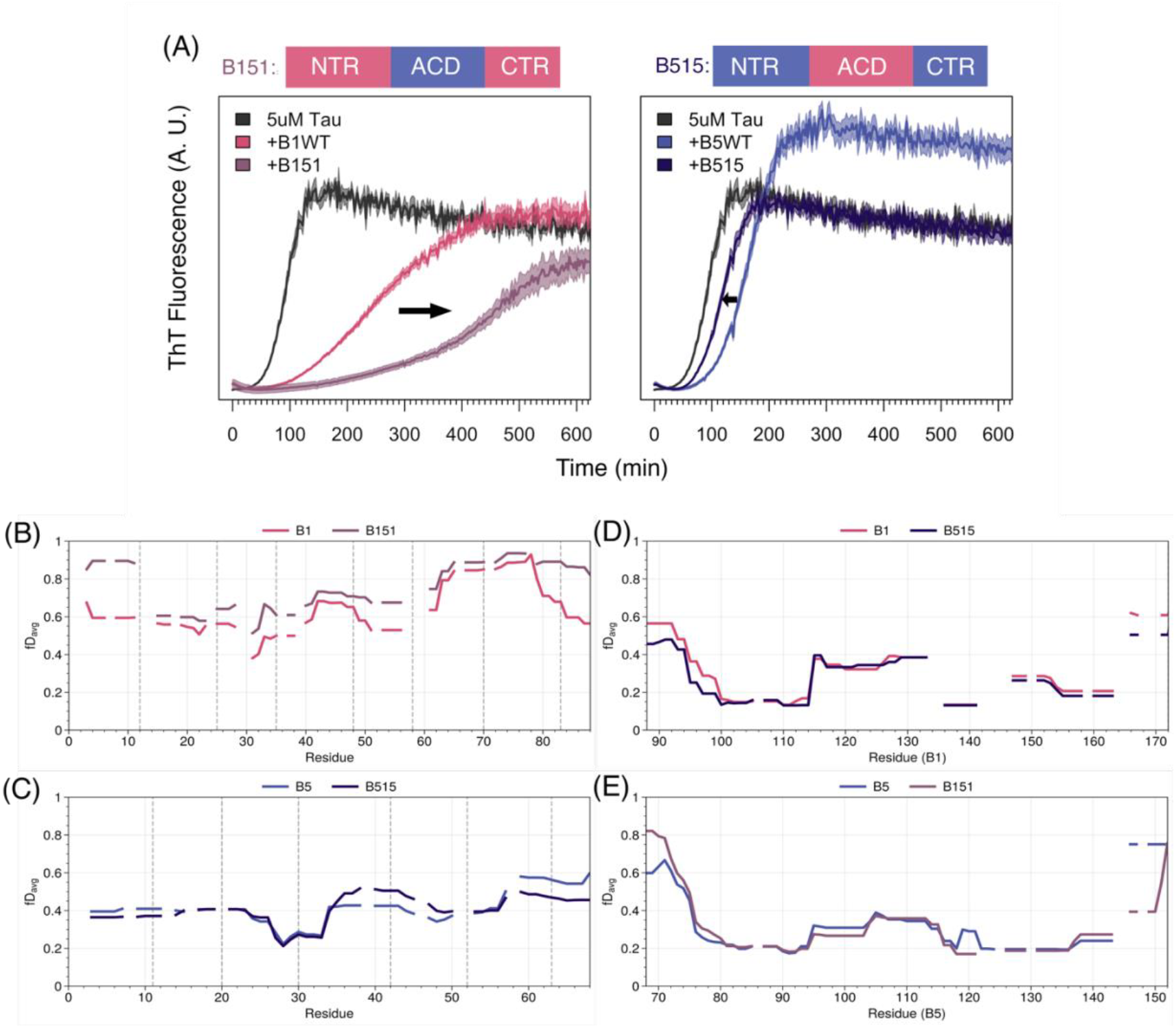
Exchanging the ACDs has opposite effects on HSPB1 and HSPB5 chaperone activities. (A) In vitro tau aggregation assay using ThT comparing WT activities of HSPB1 (Left) and HSPB5 to their ACD-swapped constructs (Right). Aggregation reactions are performed at pH 7.5, 37 °C using 5 μM Tau and 0.25 molar equivalent of small heat shock protein. Both fibrilization reactions are induced with 1 mg/mL polyphosphate. Black arrows show direction of changes in activities going from WT to ACD-swapped construct. (B-C): Comparison of average deuteration uptake for the NTRs of HSPB1 and B151 (B) and HSPB5 and B515 (C). Average deuteration level calculated based on unimodal analysis is shown for corresponding residues. Figure 2B shows HSPB1-WT (pink) compared to B151 (purple) after 4 seconds incubation in deuterated buffer. Figure 2C shows equivalent for HSPB5-WT (Blue) and B515 (Dark blue). (D-E):ACD-matched HDX-MS comparison of HSPB1 and HSPB5 WT vs ACD-swapped constructs. Figure 2D shows HSPB1-WT ACD (pink) compared to B515 ACD (Dark blue). Figure 2E shows equivalent for HSPB5-WT (Blue) and B151 ACD (purple).

#### HDX-MS reveals altered NTR scaffolding in chimeras

To directly assess how ACDs identity influences NTR sequestration, we compared WT and chimeric constructs by single time point hydrogen-deuterium exchange mass spectrometry (HDX-MS). HDX-MS uniquely enables near-residue-level interrogation of quasi-ordered polydisperse oligomers of sHSPs in their native state. Because NTR sequestration is most evident at very short exchange times, we performed analysis following a 4-second incubation in D_2_O-based buffer.

Peptide-level data were aggregated to obtain near-residue-level fractional D-update values, using the approach described in (Tuttle 2025), shown in Figure 2. To ensure comparison between identical sequences, data are plotted by region: NTR comparisons include WT- HSPB1 versus B151 and WT-HSPB5 versus B515 (Figure 2B); ACD comparisons include WT- HSPB1 versus B515 and WT-HSPB5 versus B151 (Figure 2C). Additional comparisons of overall exposure are provided in Figure S2.

Across all constructs, the ACD is the most protected region, displaying the lowest fractional D-uptake values. The only substantial perturbations within the ACD observed across constructs are near the NTR-ACD boundaries (84-100 for B1, 64-80 for B5): this region is more exposed in B151 compared to WT-HSPB5 (Figure 2C) and more protected in B515 compared to WT-HSPB1 (Figure 2D). The affected residues include those that form the β3-strand of the ACD β-sandwich (PDB: 4MJH, PDB: 2N0K). In some solved ACD structures, an additional β-strand (“β2”) formed by residues at the C-terminal end of the NTR pairs with β3-strand (Hochberg 2014; Rajagopal 2015). Reduced protection of β3 in B151 suggests an incompatibility between the β2-forming sequence of HSPB1 and the β3 strand of HSPB5. Conversely, increased protection in B515 suggests that some region of the HSPB5 NTR interacts with the β3-strand of HSPB1 ACD.

#### CTRs exposure is insensitive to ACD identity

The CTRs in HSPB1 and HSPB5 display high deuterium uptake at 4 seconds, consistent with their known disorder and accessibility in oligomers. Coverage across the CTRs was insufficient to allow more detailed analysis due to limited redundancy and resolution (SI Figure 3).

**Figure 3.**
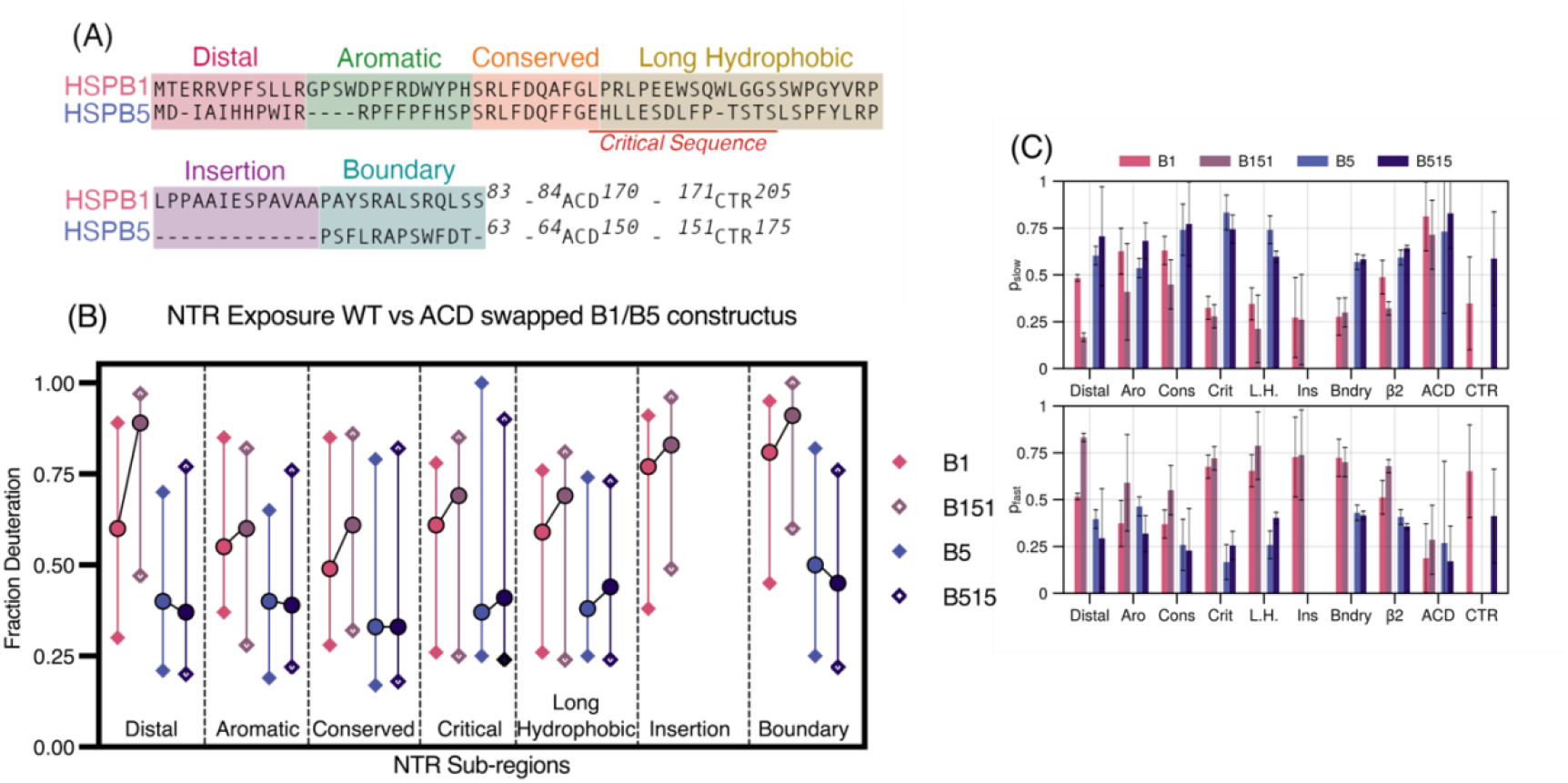
Comparison of the NTRs by subregion. (A) Sequence alignment of HSPB1 and HSPB5 NTR residues. The NTRs of HSPB1 and HSPB5 are divided into 5-6 short subregions, following Clouser 2019 nomenclature for HSPB1. All peptide data was aggregated to obtain near-residue-level fractional deuterium-uptake values, followed by bimodal analysis using the pyHXExpress workflow (Tuttle 2025). Average fractional D-uptake for each NTR- subregion, calculated from the centroids of unimodal fits, is shown as circles (○). Deuteration levels of the two populations (slow and fast states) determined from bimodal analysis are plotted as diamonds (◊) in the corresponding color scheme. (C) Relative populations of the slow (*p*_*slow*_) and fast (*p*_*fast*_) exchange modes for each NTR-subregion, as well as for the ACD and CTR are shown.

#### NTR accessibility correlates with chaperone activity

Comparison of NTRs in their native context shows that HSPB1 NTR is substantially more solvent-exposed than HSPB5 NTR under identical conditions, with average fractional D-uptake values exceeding 0.5 for HSPB1 and below 0.5 for HSPB5 (compare the lighter traces in Figure 2B and 2C). ACD identity strongly modulates HSPB1 NTR exposure: in the B151chimera, the HSPB1 NTR exhibits increased fractional D-uptake, consistent with the NTR being less effectively sequestered by the ACD of HSPB5 as inferred from the chaperone assay data.

In contrast, HSPB5 NTR is comparatively less sensitive to ACD swapping. In the B515 chimera, we observe only modest and localized changes in fractional D-uptake values, including a small increase in protection of residues ∼60-68 and decreased protection in the vicinity of residues 35-48. Notably, B151, the chimera with the most prominent chaperone activity, displays the highest overall NTR exposure, and B515, which displays slightly lower chaperone activity than the WT-HSPB5, has a modest decrease in D-uptake primarily in a single region, the “Boundary” region that links the NTR to the ACD. (Figure 2C and Figure 3B).

### Subregion-specific differences in NTR sequestration between HSPB1 and HSPB5

Previous studies have revealed that NTRs are organized as a series of short subregions with different properties (Figure 3A). The Distal, Conserved, and Boundary subregions contribute to the quasi-ordered organization of oligomers through inter-subunit interactions with ACDs and the Long Hydrophobic subregion is implicated in modulating chaperone activity against γD-crystallin. Thus, NTR subregions can serve both structural roles (via sequestration within the oligomer) and provide chaperone function directly. HDX-MS provides a direct readout of the reorganization within oligomers that arises from differences in how each ACD variant sequesters the NTR. Here we analyze previously defined NTR subregions (Figure 3A) in WT and ACD-swapped constructs.

HSPB1 and HSPB5 oligomers exhibit bimodal HDX behavior across their NTRs, reflecting conformational heterogeneity within quasi-ordered regions (Clouser 2019; Woods 2025). Bimodal exchange indicates the presence of at least two distinct populations—more protected (slower exchanging) and more solvent-exposed (faster exchanging)—that correspond to sequestered versus exposed states. Across all four constructs, most NTR subregions display bimodal behavior, consistent with earlier reports. For each subregion, we extracted: 1) fractional D-uptake values for the slow- and fast-exchanging modes (diamond markers in Figure 3B), and 2) the relative populations of these modes (*p*_slow_ and *p*_fast_, where (*p*_slow_ + *p*_fast_ = 1.0)). Average D-uptake values from unimodal fits are also presented in Figure 3B and SI Figure 3.

Overall, the most profound differences observed between a WT oligomer and its ACD- swapped counterpart are for HSPB1 and B151. In B151, multiple NTR subregions show substantial increases in the population of fast-exchanging species, indicating increased numbers of exposed regions in the oligomeric ensemble. As well, the slower-exchanging population of the Distal and Boundary regions, known to be involved in key NTR-ACD interactions, show increased fractional D-uptake values. Hence, when the HSPB1 NTR is scaffolded by the HSPB5 ACD, not only are exposed conformations more highly populated, but the slower-exchanging population is less well-sequestered. By comparison, the HSPB5 NTR is much less sensitive to the ACD identity, with B515 exhibiting only subtle changes in bimodal distribution metrics.

#### Distal and Boundary regions are most sensitive to ACD identity

The Distal region displays the largest ACD-dependent effects. In B151, the fractional D-uptake value of the slower-exchanging population increases from ∼0.3 to ∼0.47 (Figure 3A) accompanied by a smaller increase in uptake for the fast population. Partitioning between the two states shifts dramatically, from approximately equal populations in WT-HSPB1 to a majority of the Distal region occupying solvent-exposed states in B151. Thus, a larger fraction of the HSPB1 Distal subregion is exposed in the context of HDPB5’s ACD and the protected population is less well sequestered.

A similar trend is observed for the Boundary region at the C-terminal end of the NTR. In HSPB1 constructs, the unimodal fractional D-uptake increases from 0.81 to 0.91 upon ACD swap. In contrast, it decreases from 0.5 to 0.45 when the ACD is swapped in HSPB5. The differences in HSPB1’s Distal subregion arise primarily from increased D-uptake of the slower-exchanging population, rather than a change in the partitioning between populations, indicating that the slower-exchanging state of the HSPB1 Boundary region is less well sequestered by the HSPB5 ACD.

Both the Distal and Boundary regions mediate key NTR–ACD interactions. The Distal region contains an “IxI” motif that inserts into the ACD Edge groove: HSPB5 carries a canonical ^3^IAI motif and HSPB1 contains a non-canonical sequence of alternating hydrophobic residues, ^6^VPFSL. The HDX results imply that this non-canonical motif is not well accommodated by the HSPB5 Edge groove. In contrast, the more conventional ^3^IAI motif in HSPB5 is effectively sequestered by the HSPB1 ACD.

The Boundary region can form a β2 strand that aligns antiparallel to the β3 strand of the ACD at the dimer interface. Upon ACD swapping, deuteration patterns diverge: the Boundary region becomes more exposed in B151 but modestly gains protection in B515. While it is difficult to rationalize the behavior in specific structural terms, we note that the HDX trends of higher/lower Boundary region exposure correlate with higher/lower tau chaperone activity.

#### Aromatic and conserved regions respond differently to ACD swapping

The Aromatic subregions which to date has no defined function, are affected by ACD identity but respond in distinct ways. In B151, the population of fast-exchanging peptides increases, while the slow mode takes up deuterium even more slowly. In B515, the population of exposed Aromatic region is decreased, while the fast-exchanging population takes up deuterium faster. Thus, a larger fraction of Aromatic regions is exposed in B151 than in WT HSPB1 oligomers and a smaller fraction is exposed in B515 than in WT HSPB5.

In both WT oligomers, the Conserved region which binds to the ACD Central (dimer interface) groove is the most protected region of the NTR and a majority of conserved regions are sequestered (*p*_slow_ > 0.5). In B151, the partitioning is altered, with more than half populating more exposed states, indicating less effective sequestration of the HSPB1 Conserved region by HSPB5 ACD.

#### HSPB1-specific Insertion subregion is largely insensitive to ACD identity

The Insertion region, unique to HSPB1, is mostly exposed, with only 26% sequestered. While its partitioning does not change in the context of the HSPB5 ACD, the sequestered population has a higher fractional D-uptake value, indicating it is less accommodated by the heterologous ACD.

### Contributions of HSPB1 and HSPB5 NTR subregions in the delay of tau aggregation

The subregion-specific differences in NTR exposure observed by HDX-MS prompted us to ask whether exposure of particular NTR elements correlates with the ability to delay tau aggregation. To identify NTR subregions that contribute to tau chaperone activity, we generated a panel of subregion chimeras between HSPB1 and HSPB5 by swapping a single previously defined NTR subregion (Figure 3A). For each chimera, tau chaperone activity and oligomer size distributions were measured at pH 7.5 and pH 6.5. Chaperone activity is reported as the fold-change in T_50_ relative to T_50_ for tau alone (“T_50_ fold-change”), with values for WT and the ACD-swaps, B151 and B515, included as reference points (Figure 4A). This approach allowed us to assess the contributions of individual NTR elements within the oligomeric contexts imposed by each sHSP.

**Figure 4.**
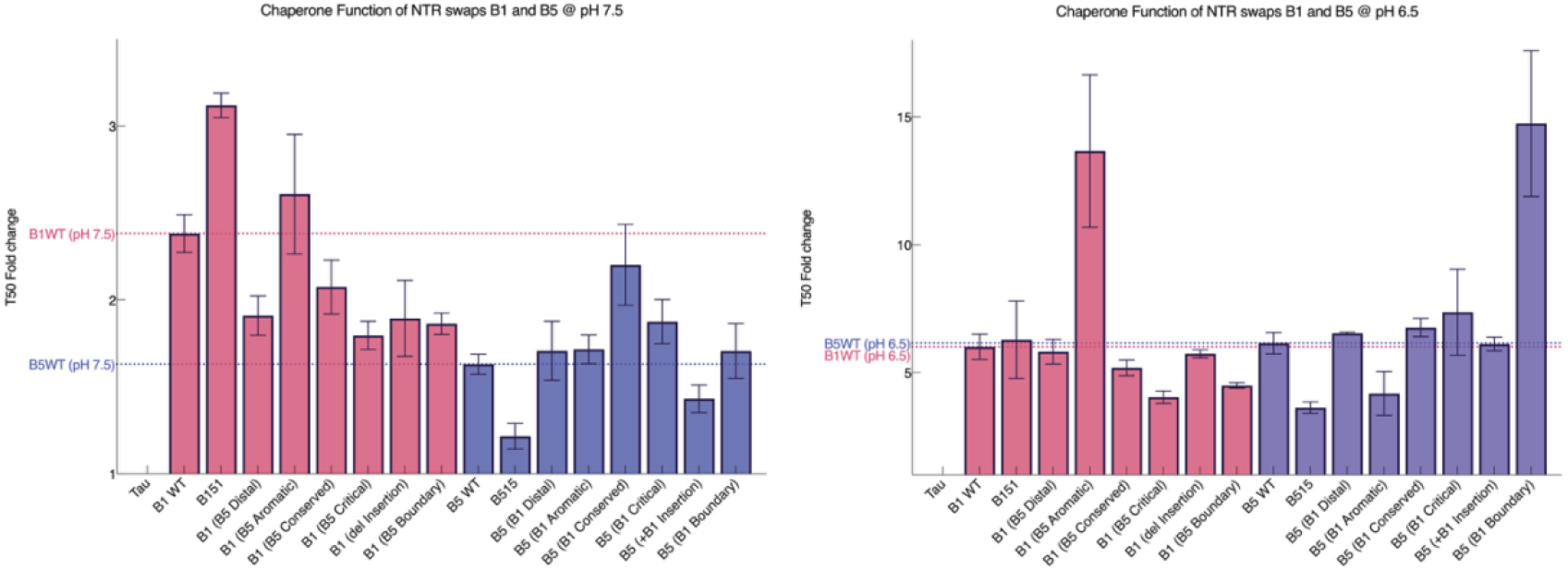
Effects of swapping individual subregions between HSPB1 and HSPB5. (A) Effects of swapping individual NTR sub-regions between HSPB1 and HSPB5 on tau chaperone activity. T_50_ fold change values for NTR sub-region swapped constructs of HSPB1 and HSPB5 constructs for assays performed at pH 7.5 (Left) and at pH 6.5 (Right). Bar graphs represent the average T_50_ values, with dotted lines representing the average T_50_ of HSPB1 and HSPB5 wild-types. T_50_ values for samples containing HSPB1 or HSPB5 were normalized to T_50_ of tau-only controls from the corresponding assays. Error bars indicate the SEM of T_50_ values calculated from 2-3 independent assays, with each assay containing 4-6 replicates.

#### Subregion contributions under basal conditions

At pH 7.5, WT-HSPB1 is an effective chaperone and B151 is further activated. Among the subregion swaps in the HSPB1 context, the chimera containing the Aromatic region of HSPB5 (“HSPB1-(HSPB5 Aromatic)”) uniquely exhibited enhanced activity relative to WT-HSPB1. All other NTR subregion swaps resulted in reduced activity, yielding T_50_ fold-change lower than WT- HSPB1.

At pH 7.5, WT-HSPB5 has only modest activity and swapping the ACD (B515) further decreases activity. Among the NTR subregion swaps, both the Conserved and Critical regions enhance activity, although neither restored activity to the level of HSPB1. Notably, the Conserved regions of HSPB5 and HSPB1 differ by only two amino acid differences, yet this minimal difference is sufficient to tune chaperone activity. This parallels previous findings in which three amino-acid substitutions distinguish the chaperone activities of HSPB5 and lens-specific sHSP, HSPB4 towards a lens client protein, *γ*D-crystallin (Woods 2023).

#### Subregion contributions under activating conditions

At pH 6.5, both wild-type HSPB1 and HSPB5 are activated and exhibit comparable chaperone function, providing a baseline for evaluating the effects of subregion swaps. Despite already being in activated states, both proteins are further activated by specific swaps (Figure 4). The most pronounced effects arise from the Aromatic and Boundary regions, followed by the Critical and Conserved regions, with only weak or no effects in the Distal or Insertion region.

#### Aromatic and Boundary regions dominate tau chaperoning activity

The Aromatic and Boundary swaps produce the strongest effects and these are reciprocal between the two analogous chimeras. HSPB1-(HSPB5 Aromatic) shows enhanced activity, while HSPB5- (HSPB1 Aromatic) has reduced activity. These results indicate that HSPB5’s Aromatic region carries intrinsic tau chaperone activity that can be transferred across oligomeric contexts. Consistent with this interpretation, the Aromatic swap is the only one that decreases HSPB5 activity under conditions where HPSB5 is activated. Furthermore, installation of the Aromatic region of HSPB5 into HSPB1 greatly enhances its activity. Altogether, the observations strongly point to a direct participation of the HSPB5 Aromatic subregion in tau chaperoning function.

A similar reciprocal pattern is observed for the Boundary subregion. HSPB5-(HSPB1 Boundary) has greatly enhanced activity, whereas HSPB1-(HSPB5 Boundary) has slightly reduced activity. Together with the HDX-MS data (above), these findings suggest that the HSPB1 boundary region, particularly when less sequestered, contributes directly to tau chaperoning.

Swapping either the Critical or Conserved regions also shift activity in opposite directions, but with only modest effects. In contrast, Distal region swaps have no measurable effect, despite the large ACD-dependent effects observed by HDX at basal condition. The lack of direct involvement of the Distal subregion in tau chaperoning is consistent with previous observations (Baughman 2020). The results show that it is not exposure of a subregion *per se* that leads to chaperone activity and that the nature/identity of the subregion matters.

### Oligomeric ensemble sizes of HSPB1 and HSPB5 constructs do not directly correlate with chaperone activities

sHSPs oligomer size distributions are sensitive to environmental conditions, post-translational modifications, and mutations (Jehle 2011; Peschek 2013; Berkeley 2025) and have been proposed to influence chaperone function. To investigate possible correlations between chaperone function and oligomer size, we compared the molecular weight distributions of WT and chimeric oligomers using mass photometry.

#### Baseline oligomeric distributions at pH 7.5

At pH 7.5, where HSPB1 is the more active chaperone, HSPB1 oligomers are markedly more polydisperse than those of HSPB5 (Figure 5A & B). HSPB1 displays a bimodal distribution of large oligomers (Average molecular weight 406 kDa or ∼20mers) and smaller species including tetramers. In contrast, HSPB5 forms a single, relatively narrow distribution centered around 443 kDa, with no detectable species below ∼250 kDa. These differences suggest weaker inter-subunit interactions within HSPB1 homo-oligomers compared to those of HSPB5.

**Figure 5.**
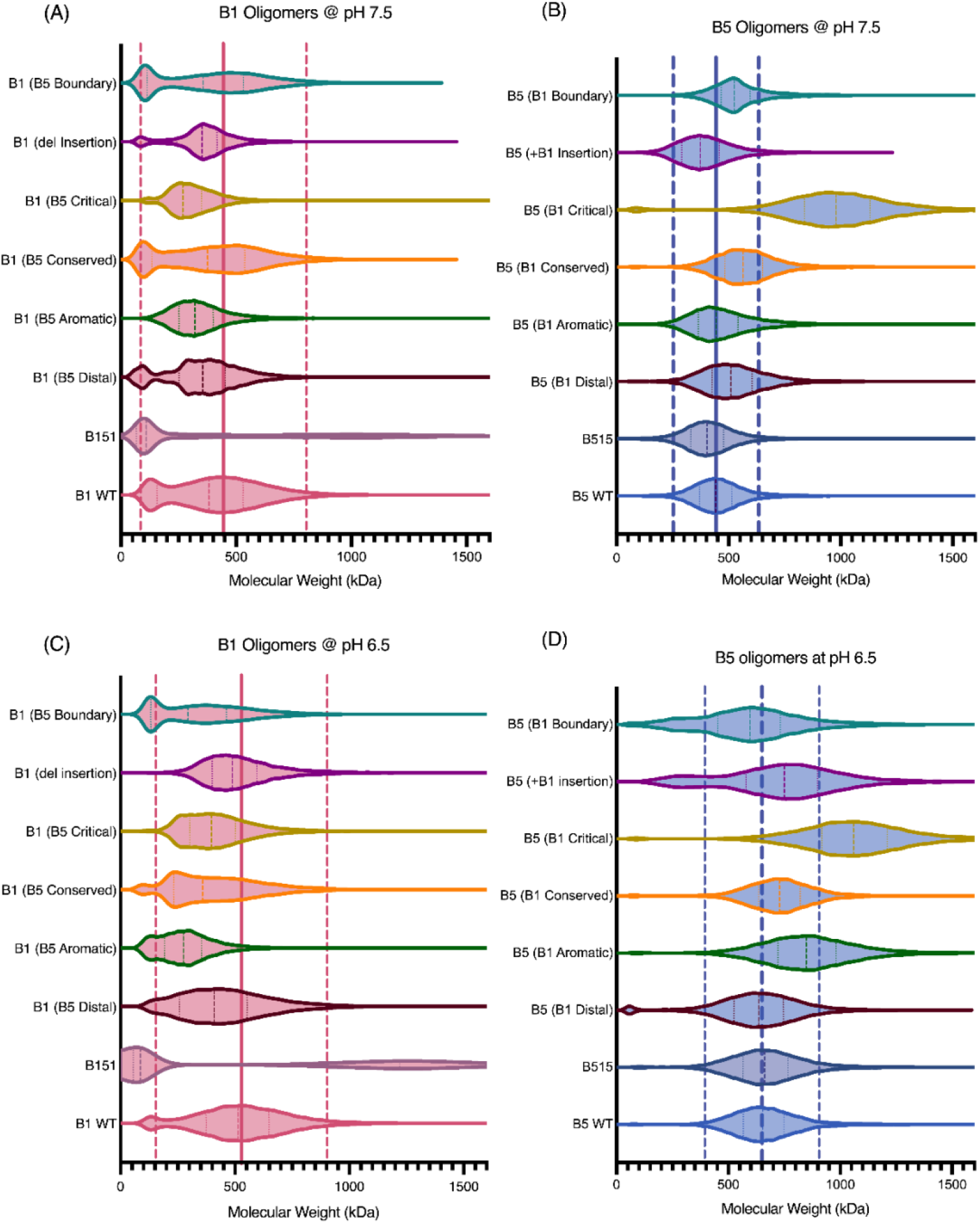
Effects of subregion swaps on oligomeric size. The molecular weight distributions of HSPB1 and HSPB5 constructs are measured by Mass Photometry. Each measurement is conducted at 1 μM final total concentration of sHSP using dilution-free measurements. The samples are incubated for 3 hours at 37 °C at the final concentration in PBS under reducing condition at the appropriate pH. The mass photometer is calibrated using beta-amylase prior to the measurements. Solid line is drawn at the calculated average molecular weight of the large oligomers, with the dashed lines placed at ± 2 standard deviations from the mean.

#### Effects of NTR subregion swaps at pH 7.5

To determine whether NTR swaps influence oligomerization, we examined the size distributions of all NTR-swap chimeras at pH 7.5. HSPB5 constructs were largely insensitive to NTR subregion swaps, with the exception of the Critical subregion swap, which results in a pronounced increase in ensemble size (WT = 443kDa vs. critical swap = 978kDa). In contrast, HSPB1 constructs were highly sensitive, with the relative populations of the smaller and larger-sized species changing in virtually every case. Aromatic and Critical swaps increase the population of larger oligomers, while Distal, Conserved, and Boundary swaps favor smaller species. Notably, replacing HSPB1’s Aromatic region with that of HSPB5 yielded a unimodal distribution resembling that of WT- HSPB5.

#### pH-dependent effects at pH 6.5

Both WT oligomers display a pH-dependent increase in oligomeric size, forming larger species at pH 6.5. Under these activating conditions, HSPB5 constructs become more sensitive to changes within the NTR. Critical and Aromatic swaps shift distributions to higher molecular weight, while Boundary and Insertion swaps favor smaller oligomeric species. Notably, introduction of the unique Insertion region or the Boundary region of HSPB1 into HSPB5 broadens the size distributions, producing profiles more characteristic of HSPB1 oligomers. In contrast, HSPB1 constructs at pH 6.5 show changes in both ensemble shape and molecular weight distribution, with no consistent trend across swaps.

#### Lack of correlation between oligomeric size and activity

Several prior studies have proposed a relationship between sHSP oligomer size and chaperone activity (Peschek 2013; Jovcevski 2015). While isolated observations in our dataset could be interpreted as supporting such a correlation—for example B151 exhibits both enhanced activity and smaller oligomers relative to WT-HSPB1—analysis across the full dataset of constructs and conditions reveals no consistent relationship between oligomer size or polydispersity and tau chaperone activity.

## Discussion

Our results demonstrate that HSPB1 and HSPB5 suppress tau aggregation through distinct, sequence-encoded strategies that are implemented by differential exposure of their N-terminal regions (NTRs). This divergence reflects a broader principle of sHSP function: chaperone activity is governed not simply by the presence of functional domains, but by how intrinsically disordered regions are scaffolded, sequestered, and released within dynamic oligomeric assemblies.

### Division of labor between HSPB1 and HSPB5

HSPB1 suppresses tau aggregation under basal conditions, whereas HSPB5 requires stress-induced activation such as lowered pH to become effective. As both proteins are constitutively expressed in neurons, this behavior suggests a functional division of labor. HSPB1 may provide continuous, long-term protection against tau misfolding, while HSPB5 functions as a rapidly deployable reserve that is mobilized under acute stress. Upon activation, HSPB5 becomes as effective as HSPB1, transiently boosting cellular protection when proteotoxic stress escalates.

### Sequence-encoded NTR accessibility

These functional differences arise largely from intrinsic properties of the NTRs. HSPB5’s NTR is more hydrophobic and histidine-rich, favoring sequestration at neutral pH and conferring sensitivity across a physiologically relevant pH range. In contrast, HSPB1’s more polar NTR is less sequestered under basal conditions. Importantly, these properties are retained in NTR–ACD chimeras, demonstrating that solvent accessibility is encoded primarily in NTR sequence rather than imposed by ACD identity.

### NTR–ACD interactions tune activation

The observations from the ACD-swaps can be rationalized in terms of known NTR-ACD intra-subunit interactions that can tune NTR accessibility and activity.

The Distal region of HSPB1, containing a non-canonical “IxI” motif (^6^VPFSL) shows balanced exposure in its native context but becomes predominantly exposed when paired with the HSPB5 ACD. This suggests that the HSPB1 ACD edge groove has evolved to accommodate its unusual Distal “IxI” motif, whereas the HSPB5 ACD does not. On the other hand, the canonical IxI motif in the HSPB5 Distal sub-region (^3^IAI) is efficiently sequestered by both ACD edge grooves. This difference likely promotes greater NTR release and thus activity in HSPB1 under basal conditions, while HSPB5’s knob-into-hole IxI interaction enables sequestration until stress triggers activation. Notably, the same ACD grooves that bind NTR motifs also engage co-chaperones and clients, including tau, pointing to shared mechanisms for activation.

A second regulatory layer involves the Boundary region, which can form an additional β- strand (β2) by pairing with the ACD β3 strand. Disrupted pairing in chimeras correlates with increased NTR exposure and enhanced activity, suggesting that HSPB1 and HSPB5 NTR-ACD boundary may exist in different conformations at basal state: HSPB1 NTR boundary sub-region is mostly pointing away from the ACD, thereby enhancing overall NTR exposure, whereas HSPB5’s β2 strand patches with the ACD’s β3 strand, thus decreasing overall exposure of HSPB5 NTRs. Such subtle changes in NTR-ACD interactions can tune exposure within the oligomers, and thus activity of HSPB1 and HSPB5.

### Modular determinants of tau specificity

Sub-region swaps reveal that tau chaperone activity is not evenly distributed across the NTR but rather is concentrated in specific subregions. Notably, installation of the Aromatic region of HSPB5 (^12^RPFFPFHSP^20^) into HSPB1 greatly enhanced tau chaperone activity while replacement of that sequence with the HSPB1 Aromatic sequence led to a loss of chaperone activity in HSPB5 (Figure 4). Installation of the HSPB1 Boundary region (^71^PAYSRALSRQLSS^83^) into HSPB5 led to a similar enhancement of its activity and the reciprocal swap led to decreased activity. These effects are most apparent under activating conditions at pH 6.5, when NTRs are more exposed.

Although these regions differ substantially in sequence composition, both may function either as direct tau-binding sites or as regulatory hubs that integrate environmental signals. The Aromatic region of HSPB5 has a His residue that could enable pH-dependent activation and both the Aromatic and Boundary regions contain the phosphorylation sites that activate HSPB1 and HSPB5 (S15, S78, S82 in HSPB1; S19, S59 in HSPB5). Notably, across the ten human sHSPs, phosphorylation sites cluster in the Boundary and Aromatic regions, suggesting that the positions and properties of these subregions within otherwise variable NTRs have conserved modulatory mechanisms.

Our findings support a modular model in which different NTR sub-regions confer specificity toward distinct aggregation pathways. The Aromatic region of HSPB5 governs tau aggregation suppression, whereas the Critical region of HSPB5 drives protection of γD- crystallin aggregation in the eye lens (Woods 2023). Tau and γD-crystallin undergo different aggregation pathways, forming fibrils and amorphous aggregates, respectively. We propose that the ability to engage different types of clients and their early-aggregating species may therefore arise from a diversity of sequences (subregions) within sHSP NTRs that can be reconfigured to meet a variety of cellular conditions and clients.

### Disease relevance

This framework also provides insight into disease mutations. Prior HDX-MS studies of HSPB1 disease variants show altered exposure across multiple NTR sub-regions, with a shared increase in Boundary region accessibility (Clouser 2019). These mutations may therefore dysregulate a key structural control point that links oligomer architecture to chaperone output. The formation of hetero-oligomers between HSPB1 and HSPB5 (Aquilina 2013; Mymrikov 2020; Shatov 2020) further suggests that cells may dynamically tune NTR accessibility in response to client load or stress.

## Conclusions

In summary, our data define a mechanism in which sHSP chaperone activity emerges from dynamic, modular exposure of NTR sub-regions embedded within a quasi-ordered oligomeric network. Sequence-encoded stress sensors and ACD-mediated scaffolding jointly determine which regions are exposed, when, and to what extent. This model explains how sHSPs achieve both client specificity and environmental responsiveness, and why certain clients may be preferentially protected under defined cellular conditions. More broadly, it identifies NTR–ACD interactions and NTR sub-regions as promising targets for therapeutic strategies aimed at modulating sHSP function in neurodegenerative disease.

## Materials and methods

### Protein expression and purification

Tau4RD was expressed in *Escherichia coli* BL21 DE3 cells (New England BioLabs (NEB)). The cells were grown at 37 °C until reaching an OD_600_ of 0.6-0.8, then induced overnight with 1 mM IPTG with the temperature adjusted to 16 °C. Cells were harvested by centrifugation and resuspended in 50 mM Tris-base, 500 mM NaCl, 10 mM Imidazole (Buffer A). Cells were lysed by a homogenizer after adding DNase, RNase, PMSF, and HIS- protease inhibitor. The lysed cells were centrifuged then the lysate was filtered (0.45 μM) and applied to HisTrap column equilibrated in Buffer A. Bound proteins were eluted by Nickel elution buffer (Buffer A + 250 mM Imidazole). HIS-TEV protease was added to elutant and dialyzed overnight in Buffer A with 2 mM DTT added. Dialyzed protein was reapplied to HisTrap column to collect the protein without the His-tag to be concentrated to ∼6 mL. The concentrated protein was injected 2mL at a time on sdx200 column in final buffer containing 25 mM NaPi, 100 mM NaCl, 0.5 mM EDTA, 2 mM. Fractions containing the protein was pooled and concentrated with a 3,000 kDa Molecular Weight Cut Off (MWCO) Amicon Ultra Centrifugal Filter.

HSPB1 and HSPB5 constructs were expressed in *Escherichia coli* BL21 DE3 cells (New England BioLabs (NEB)). The cells were grown at 37 °C until reaching an OD_600_ of 0.6-0.8, then induced overnight with 500 μM IPTG with the temperature adjusted to 22 °C. Cells were harvested by centrifugation and resuspended in lysis buffer (20 mM Tris-base, 100 mM NaCl, 1 mM EDTA, pH 7.5). Cells were lysed by a homogenizer in presence of DNase, RNase, lysozyme, and protease-inhibitor. HSPB1 purification was performed by Ni-NTA, following the tau purification protocol, but the tag cleavage was performed with SENP1. Following dialysis and tag-cleavage, HSPB1 sample was desalted into 20mM Tris Buffer at pH 8.0 using a 5mL HiTrap Desalting column, then loaded on to a 5 mL HiTrap Q High Performance column and eluted over a 100 mL gradient into the same buffer containing 1 M NaCl. The sample was then concentrated using a 10,000 kDa MWCO Amicon Ultra Centrifugal Filter before size exclusion chromatography. HSPB5 purification protocol was followed as described previously (Woods 2022).

### Fibril formation assays

Fibrilization assay of Tau was performed on a low-binding 96-well plate using a CLARIOstar microplate reader. Each reaction contained 200 μL of sample with 5 μM Tau4RD, 1 mg/mL polyphosphate, 130 μM Thioflavin T (ThT), and 1.25 μM of chaperones in 25 mM NaPi, 20 mM NaCl, 0.5 mM EDTA, and 5 mM DTT buffer. The plate reader was shaken for 1 minute prior to fluorescent measurement at 440 nm excitation with 485 nm emission every 2.5 minutes. The enhanced dynamic range setting was applied to sample measurements. Each condition was performed in 4-6 replicates. Samples containing either HSPB1 or HSPB5 were pre-incubated in buffer and ThT for 3 hours at the same assay setting to let the oligomers and ThT come to an equilibrium on the plate reader. Tau4RD and polyphosphate were added after 3 hours to initiate the aggregation reaction.

### Hydrogen-deuterium exchange mass spectrometry

All samples were incubated at 37 °C for 3 hours in water-based buffer and then cooled to room temperature before undergoing deuterium exchange. HSPB1 and HSPB5 constructs (0.04 mg/mL) were incubated at room temperature in 85% D_2_O-based PBS pH 7.5 buffer for 4 seconds. Fully deuterated samples were made by incubating protein in 85% D_2_O + 6 M guanidium-hydrochloride buffer at 90 °C for 30 mins. Exchange was quenched by addition of ice-cold quench buffer (1.6% formic acid) and samples were flash frozen in liquid nitrogen. Samples were automatically thawed, digested by immobilized Nepenthesin-2, and injected on a Waters Synapt G2-Si instrument using a setup built in house around the LEAP PAL system (Watson 2021). HSPB5 peptides were identified by MS/MS on a Thermo Orbitrap Fusion Tribrid instrument and MS^E^ on a Waters Synapt G2-Si followed by data analysis using ProteinProspector (UCSF) or ProteinLynx Global SERVER (Waters). The quality of peptides were assessed in HDExaminer 3.0 (Sierra Analytics). All peptide data was aggregated to achieve near-residue level resolution, and the fractional deuterium uptake was analyzed following pyHXExpress workflow for unimodal and multimodal analysis (Tuttle 2025).

### Mass Photometry

All samples were incubated at 37 °C for 3 hours in phosphate-based buffer (25 mM NaPi, 150 mM NaCl, 0.5 mM EDTA, 5 mM DTT at a pH of either 6.5 or 7.5) at a final concentration of 1 μM. Prior to sample preparation and incubation, the buffers were filtered using 3 kDa Millipore Amicon Ultracentrifugal filter. A Refeyn TwoMP mass photometer (Refeyn Ltd., UK) was used to analyze the sHSP samples. Beta-amylase was used to calibrate for molecular weights by assigning 56 kDa monomeric peak, 112 kDa dimeric peak, and 224 kDa tetrameric peaks. For sHSP measurements, 20 μL of 1 μM total protein sample was directly added to the coverslip using dilution free measurement. Each measurement was recorded over two minutes and fit to a Gaussian curve fit in Prism 10 (GraphPad Software, Boston, MA) for molecular weight determination of oligomeric species present.

## Supplementary Material (Description of “MC_SupplementaryMaterials”)

*Supplemental Figure 1:* Additional *in vitro* aggregation assays induced by heparin (A), and ACD-only controls (B). Analysis of B1 and B5 FL and ACD chaperone capacity over range of sHSP concentrations, and T_50_ analysis of ACD-swapped construct with controls.

*Supplemental Figure 2*: Full-length HDX-MS comparison of HSPB1 and HSPB5 constructs.

*Supplemental Figure 3*: Redundancy of peptides used in HDX-MS analysis

*Supplemental Figure 4*: Further analysis of HDX-MS bimodal calculations of HSPB1 and HSPB5 constructs.

## Acknowledgement

This work was supported by the National Eye Institute: 2 R01 EY017370 (R.E.K.), T32 EY07031 (M.K.J.), by the National Institute on Aging: T32 AG066574 (M.K.J.), by the National Institute of General Medical Sciences T32 GM007750 (M.C.) The authors gratefully acknowledge support from the Seattle Partnership for Research on Innovative Therapies (to A.N.) and from the Edmond H Fischer/Washington Research Fund Endowment (to R.E.K.)

