## Supplemental Information for "Function within Disorder: Small heat shock proteins use different functional regions to chaperone tau aggregation"

**Supplementary Material:**

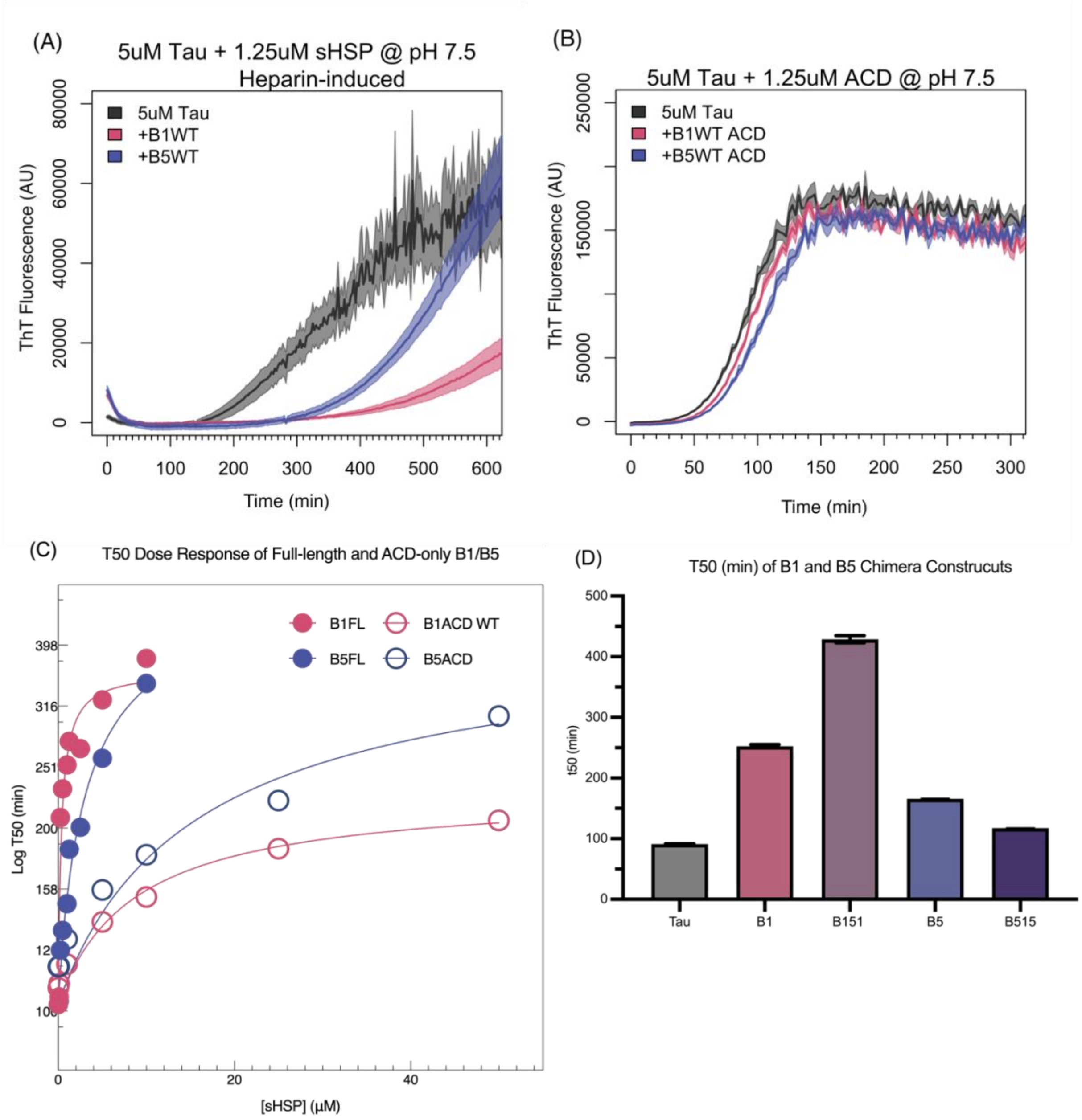


**Supplemental Figure 1**: A) HSPB1 and HSPB5 chaperone heparin-induced aggregation of tau. B) ACD-only constructs at the equivalent concentration at 1.25uM show no chaperone function. 5uM Tau aggregation is induced by 1mg/mL polyphosphate. C) T50 dose response curves for both Full-length and ACD-only constructs of HSPB1 and HSPB5, plotted on a log scale. Both Full-length constructs reach near saturation by 10uM or maximum concentration tested. HSPB5 ACD shows some titratable activity at high concentration, but the chaperone activities of both ACDs are much lower compared to equivalent concentrations of their full-length counterparts. D) Calculated T50 values of HSPB1 and HSPB5 constructs chaperone activity measured by ThT assay. SEM of 6 replicates are shown as error bars.

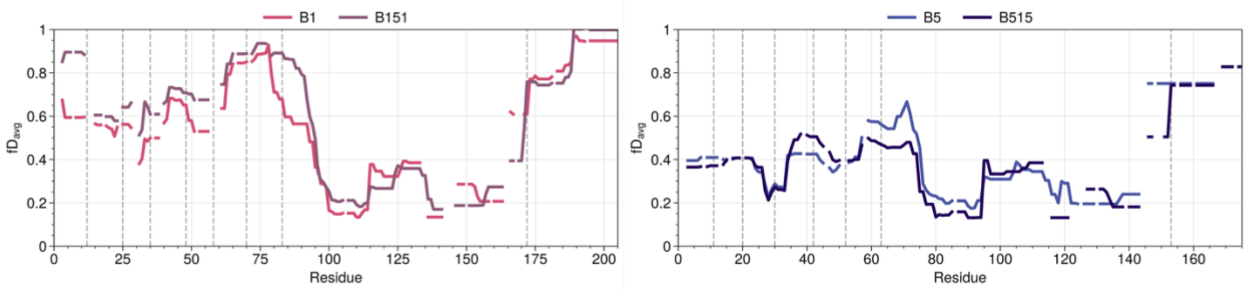


**Supplemental Figure 2**: HDX-MS full length comparison. Average deuteration level calculated based on unimodal analysis is shown for corresponding residues. Comparison of the full-length sequence for HSPB1-WT (pink) and B151 (purple) are shown in the Lef panel. Comparison of HSPB5-WT (blue) and B515 (dark blue) are shown in the Right panel. Dashed lines correspond to NTR sub-regions, ACD, and CTR residue boundaries.

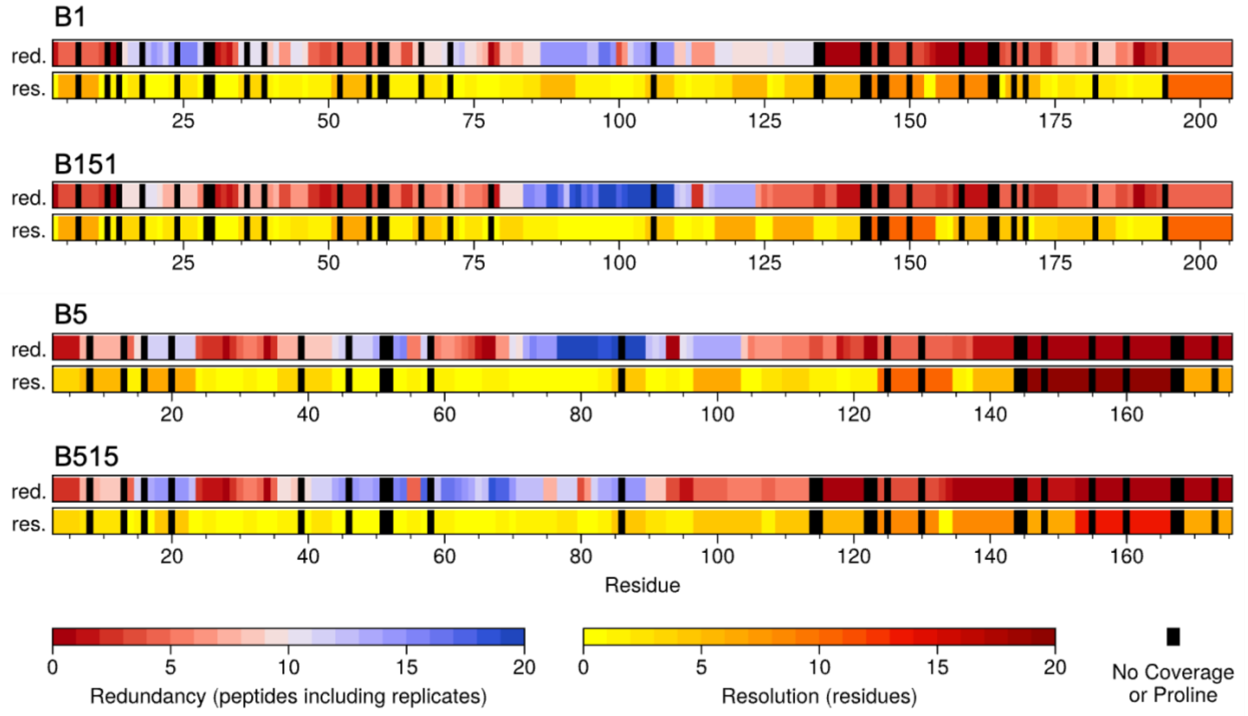


**Supplemental Figure 3**: HDX-MS Redundancy: The redundancy and resolution of peptides along the sequence are shown for HSPB1WT, HSPB151, HSPB5WT and HSPB515. Regions where there are no coverage or Proline residues depicted in black.

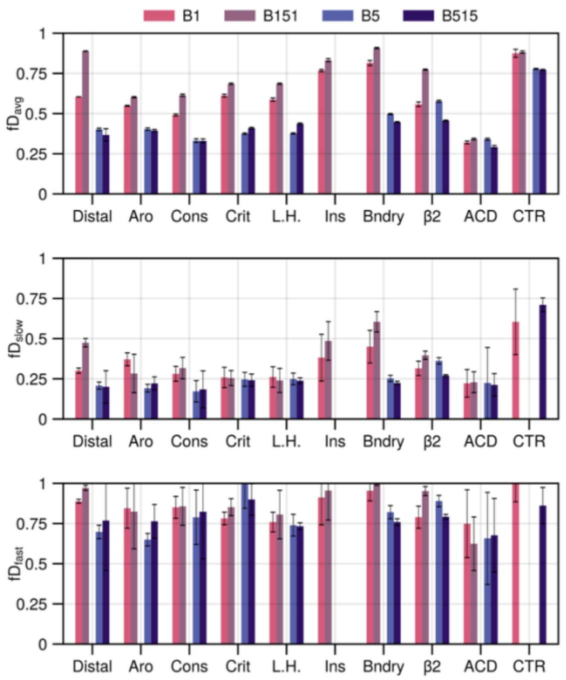


**Supplemental Figure 4**: Bimodal analysis of HSPB1 and HSPB5 constructs. Average fractional deuterium uptake for each NTR sub-regions, ACD, and CTR (Top). Fractional deuterium uptake for the slower exchanging population (middle) and fast exchanging population (bottom).
